# Chimeric U-Net – Modifying the standard U-Net towards Explainability

**DOI:** 10.1101/2022.12.01.518699

**Authors:** Kenrick Schulze, Felix Peppert, Christof Schütte, Vikram Sunkara

## Abstract

Healthcare guided by semantic segmentation has the potential to improve our quality of life through early and accurate disease detection. Convolutional Neural Networks, especially the U-Net-based architectures, are currently the state-of-the-art learningbased segmentation methods and have given unprecedented performances. However, their decision-making processes are still an active field of research. In order to reliably utilize such methods in healthcare, explainability of how the segmentation was performed is mandated. To date, explainability is studied and applied heavily in classification tasks. In this work, we propose the Chimeric U-Net, a U-Net architecture with an invertible decoder unit, that inherently brings explainability into semantic segmentation tasks. We find that having the restriction of an invertible decoder does not hinder the performance of the segmentation task. However, the invertible decoder helps to disentangle the class information in the latent space embedding and to construct meaningful saliency maps. Furthermore, we found that with a simple k-Nearest-Neighbours classifier, we could predict the Intersection over Union scores of unseen data, demonstrating that the latent space, constructed by the Chimeric U-Net, encodes an interpretable representation of the segmentation quality. Explainability is an emerging field, and in this work, we propose an alternative approach, that is, rather than building tools for explaining a generic architecture, we propose constraints on the architecture which induce explainability. With this approach, we could peer into the architecture to reveal its class correlations and local contextual dependencies, taking an insightful step towards trustworthy and reliable A.I.

Code to build and utilize the Chimeric U-Net is made available under: https://github.com/kenrickschulze/Chimeric-UNet---Half-invertible-UNet-in-Pytorch

## Introduction

Deep learning (DL) has become ubiquitous in medical diagnosis. In particular, the sub-field of *semantic segmentation* has made significant contributions to human well-being through image guided diagnosis of severe diseases (1). In semantic segmentation tasks, each pixel of an image is tasked to be classified as one of the prescribed set of classes, for example, detecting and localizing cancer in MRI scans of patients. Convolutional Neural Networks (CNNs) are the main work-horse in semantic segmentation problems, in which features are learnt from data, embracing the ability to find novel features beyond our traditional hand-crafted features. This learning based feature finding leads to CNNs being “black-boxes”, as their decision making process lacks transparency and interpretability in many applications. However, in order to prescribe DL aided healthcare to people, trust in predictions and how they were made is imperative.

In 2015, Ronneberger et al. introduced a variant on the classical encoder-decoder CNN architecture that included *skip connections* between respective blocks of the encoder-decoder, which they named the *U-Net* (2). The U-Net was originally designed for the segmentation tasks of biomedical datasets, but also found application in areas like autonomous driving (3), (4). Many modifications of the U-Net have been proposed, each driven to exploit structures in particular datasets, especially in the field of biomedical imaging (5), (6). However, *explainability* is still in its infancy for the U-Net and is an active field of research. To date, there are only a few options for Explainable A.I. (XAI) in semantic segmentation tasks (7), (8), (9).

XAI emerged out of the need to explain how and why a classification task reaches its prediction. Most XAI approaches for CNNs can be divided into two subgroups, namely perturbation-based (10), (11) and backpropagation/gradient-based approaches (12), (13), (14), (15). In practice, the gradients are computed with respect to the activations of the latent space. We can state the core XAI question in classification tasks as follows:

> (XAI *α*) – *Which region in the source contributed to that particular prediction?*

In gradient-based approaches, this question is answered by studying *saliency maps*, that is the derivative of the prediction w.r.t. the source, which highlights regions in the source, that induce strong derivatives in the prediction. In practice, those maps are, for example, superimposed directly onto the input (13) or collapsed with the target activations beforehand (12).

However, in semantic segmentation, the (XAI *α*) question needs to also incorporate spatial information into the prediction. To do this, Vinogradova et al. proposed to sum all pixel-wise class scores of interest in the prediction and evaluate the derivative of this scalar with respect to the source, which is currently the most cited XAI approach for semantic segmentation tasks (7).

Inspired by gradient-based XAI analysis, we propose a modification/constraint to incorporate the XAI motivated gradients into architectures. In particular, we present the Chimeric U-Net, which is the standard U-Net with an invertible decoder unit, i.e. a continuous invertible decoder map. This invertibility restriction on the decoder implies that the architecture:

I. has a continuous-differentiable mapping between the target space and the latent space,
II. has a simple mathematical form for the derivatives of the latent space w.r.t. to the target space and *vice versa*,
III. has the ability to remove redundant information inside the non-invertible encoder, giving a concise latent representation.

Furthermore, only considering invertibility to the latent space and not all the way back to the input space, we focus only on the features learned by the architecture, rather than the full feature space of the input.

Exploiting the invertibility of the decoder of the Chimeric U-Net, in this work, we study a novel XAI question:

> (XAI *β*) – *What is the sensitivity of the latent space w.r.t. the changes in the target prediction?*

Without loss of generality, we can think of the question (XAI *α*) as a tool for studying the sensitivity of the architecture, and (XAI *β*) as a tool to study its specificity, we will derive and explore this in the later sections of this paper.

In this work, we first demonstrate how making the decoder invertible does not hinder the network in performing semantic segmentation tasks. Then, we show how using the invertibility of the decoder enables XAI analysis, such as disentangled class information in the latent space embedding, statistical predictions of unseen data, and lastly, constructing saliency maps. The paper is structured as follows: In § we begin by introducing the Chimeric U-Net and presenting the mathematical framework for the architecture and of the subsequent XAI analysis. In § C, we present the numerical experimental setup that was used to understand the properties of the Chimeric U-Net . In § J, we present the results of how the Chimeric U-Net compares to other methods on two medical segmentation datasets. We perform XAI analysis on the medical datasets for new insights through the XAI analysis mentioned above. Lastly, in § N, we discuss the limitations, impact, and future research questions in regards to the Chimeric U-Net approach.

## Background

### A. Notation and Naming Conventions

The following are core notations of this paper: The integer product *H* × *W* denotes the height and width of the images. The integer *C* denotes the number of class/target channels. The integer *D* denotes the number of channels in the input signal/image. We use 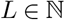 for the number of layers, we index each layer by *l* ∈ {1, …, *L*}. The operators of the architecture act on tensors, for which we use for brevity bold notations, e.g. **x**, for general tensors whose dimensions should be determined in context of its appearance. Furthermore, for a tensor **x**, we denote the *splitting*, which inherently only acts on the channel dimension of the tensor,

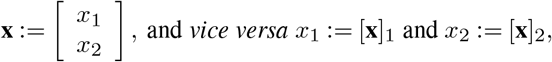

as for when we *concatenate* two tensors into a single tensor and when we want to extract the first or second split of the tensor, respectively. We propose this tensor like notation, which is notably unconventional, so that later in the article we can describe the entire decoder in a single equation. In regards to naming conventions, we interchangeably use “target” and “segmentation” for easy reading.

### B. The Architecture

We first describe the steps of the Chimeric U-Net (see Fig. 1) in plain language and proceed to reiterate the steps formally using mathematical nomenclature. To begin, the Chimeric U-Net consists of two sub-units connected by skip connections: the *non-invertible encoder* and the *invertible decoder*. The invertibility of the decoder fixes the dimensionality of the skip connections, which is determined by the number of target classes and the dimensions of the image. The encoder is non-invertible and is free to be modelled to the data of interest, with a constraint to contribute the prescribed dimensionality of the skip connections. Given the encoder is specific to the data, we state the specifics of the encoder in the numerical experiments, and from here on only focus on describing the invertible decoder.

**Fig. 1.**
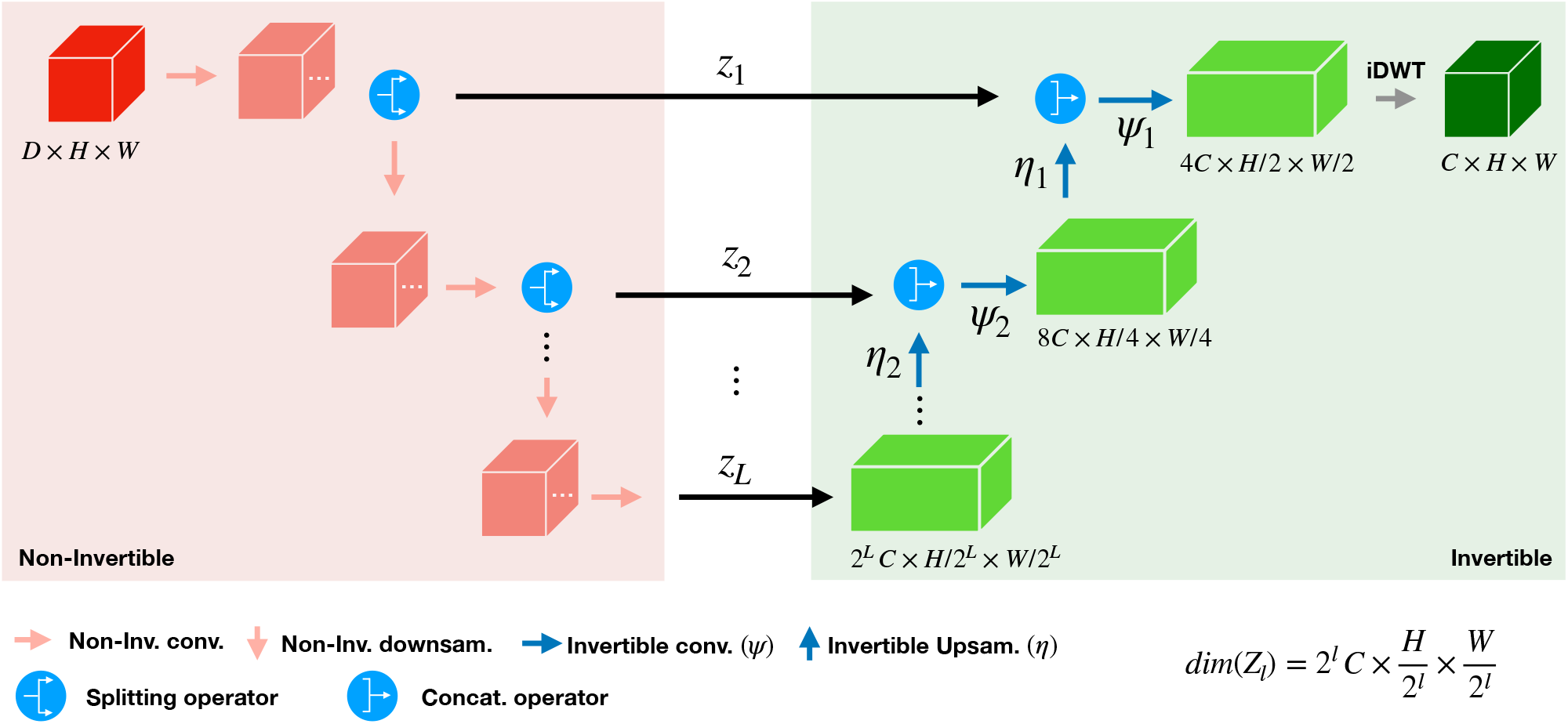
The Chimeric U-Net Schematic: non-invertible decoder connected over skip connection to an invertible decoder.

Observing the invertible decoder in Figure 1 (highlighted with green background), we see that the input to a layer in the decoder is the concatenation of the corresponding encoder layer output and the upsampled output of the layer below. The dimensions of these two tensors are the same, therefore, we concatenate them in the channel dimension and then pass it through the operations of that layer. In particular, in these middle layers, the input tensor goes through a sequence of simple *additive coupling layers* from *Normalising Flows* (16). Those additive coupling layers are in essence transforming the features in a continuous and invertible fashion. After this, the output tensor is sent to the layer above via the *invertible learnable upsampler* (17), which rearranges the pixels of the layer from, for example, *C* × *H* × *W* to *C*/4 ×2*H* × 2*C*, in order to alter the dimensionality of their input. Invertible downsampling filters are the adjoint operator of the exponential of the orthogonal skew-symmetric matrices. In that, rows when reordered into filters and convolving with an appropriate stride act as orthogonal convolutions. Since the operator is orthogonal and the involved dimensions are finite, then its inverse, i.e. the invertible upsampling filter, is just the adjoint operator. Hence, all operations in the decoder of the Chimeric U-Net are inherently invertible. This decoder layer design is that of the *Invertible U-Net* which was introduced by Etmann et al. (17).

The first and the last layers of the decoder are exceptions, that is, after the additive coupling layers of the first layer (*l* = 1) an *inverse discrete wavelet transform* (`iDWT`) is applied to the output tensor, reducing the dimensionality down to the target dimensions. Lastly, the last layer (*l* = *L*) does not receive any input from a layer below, therefore, simply passes the skip connection directly through to the layer above via the upsampler.

In summary, the decoder is rearranging-, concatenating-, and continuously deforming the points from the skip connections to the segmentation. This gives us a unique correspondence between the Cartesian product of the skip connections and the segmentation. We now present the steps of the decoder unit with mathematical nomenclature.

We begin by introducing the notation for the sets of interest. We denote the input image (*source*) set by 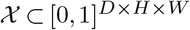, containing a stack of *D* image channels of dimension *H* × *W*. We define our target (*segmentation*) set by 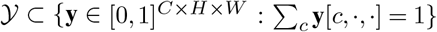, stack of *C* classes each of shape *H* × *W*, where the sum over the classes is equal to 1 for each coordinate in *H* × *W*. In addition to the input- and target sets, we have the set of skip connections, which we denote by,

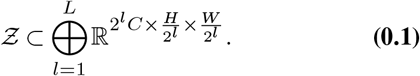

For 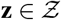 and *l* ∈ {1,…, *L*}, we denote to the *l^th^* skip connection by *z_l_*, i.e. **z** ≔ (*z*_1_,…, *z_L_*).

Using these three sets, 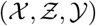, the Chimeric U-Net is defined as the mappings:

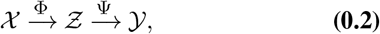

where Φ is the non-invertible encoder function and Ψ is the invertible decoder function.

We recall, the encoder function, 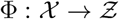, is free to be chosen to exploit the specific features of the dataset to be studied. However, the map 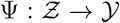 is a fixed nested function, with the evaluation of an element 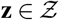 given by,

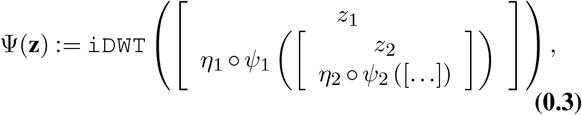

where the last term in the nesting is *z_L_*.

The invertible decoder function Ψ, Eq. (0.3), is constituted of three key functions: the inverse discrete wavelet transform, the learnable invertible upsampling operators (*η*_•_), and lastly, the normalising flow units (*ψ*_•_). In particular, for a layer *l* of the Chimeric U-Net, the learnable invertible upsampling operator is given by the map,

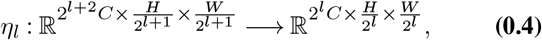

which preserves the total number of elements in the mapping from the domain to the range (17). The normalising flow of layer *l* is given by,

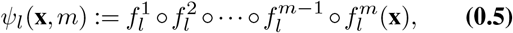

a composition of 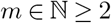 additive coupling layers of 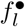, given by,

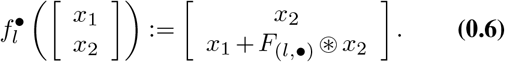

For brevity, we omit specifying *m* in the general representation of *ψ* in Eq. (0.3).

With the definitions above, we can define the inverse of Ψ, mapping points from the segmentation back into the skip connections, as follows: for 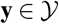,

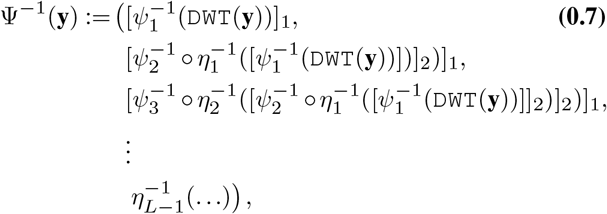

where the inverse of the normalising flows is given by,

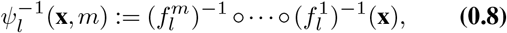

with

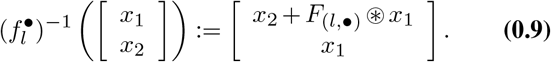

Lastly, the inverse of the learnable invertible upsampling function becomes an invertible downsampling function,

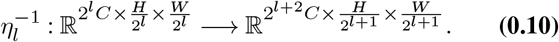

#### Remark 0.1.

*In practice, we apply a `softmax` operation after the `iDWT` to satisfy the probability constraint of the elements in the target set. This makes* Ψ *non-invertible on the whole range* 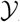, *however, we are strictly interested in the subset of the target set*, 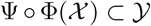, *on which* Ψ *is invertible, which here on in we denote by* 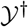. *We treat this set as the range of the architecture, arid, its latent space is the focus of this work.*

The invertibility of the decoder is a tool for bringing explainability to the U-Net. With this map, we can explain the sensitivity and the computational reasoning behind the predicted segmentation of the architecture. In the following section, we derive how the invertible decoder can be used for explainability.

### C. Latent Space

The Chimeric U-Net constructs a latent space in its skip connections, we interpret 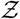 as a multiresolution latent space of the elements of 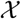. In general, studying latent spaces in imaging based tasks is an active field of research; the main focus is put on constructing meaningful latent representations of the input data for classification- and generative tasks (18). However, in the particular case of semantic segmentation tasks, the inability to untangle the multiple segmentation classes of the target in the latent space pose a challenge. That is, given the target **y** is a multi-class tensor, then each point **z** = Ψ^−1^(**y**) has entangled information of all the classes and pixels of **y**; making it hard to naturally partition 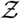 w.r.t. 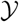.

An approach to disentangle the classes has been the use of *saliency maps* (13). In brief, saliency maps are heatmaps that highlight pixels, which contributed most to the prediction score of the target class. So far, they have been predominantly applied in image classification tasks rather than image segmentation. Probably, the most well-known of such gradient-based saliency maps is *Grad-CAM* (12) with all its recent modifications (19), (14), (20). *Grad-CAM* provides visual explanations for CNN predictions of a convolutional layer (usually the last is chosen) within the network by assigning a relevance score to each pixel. The relevance score is computed by collapsing the activations maps of the target convolutional layer, weighted by the respective gradients. In *Grad-CAM*, each channel is weighted by the same average gradient score, whereas in a more recent modification, namely HiResCAM, the *Hadamard product* of the feature maps with the respective gradients is taken before collapsing them (19). Saliency maps are an active field of research to explain Neural Network decisions. However, their evaluations, which are generally based on human interpretation (19), (21), (22), (23), are a topic of ongoing scientific debate (24), (25), (26), (27), (23).

We borrow concepts from *Grad-CAM* to help create a class-deconvolved latent space over the skip connection of the Chimeric U-Net. The invertibility and the continuity of the decoder map facilitates the computation of the *pull-back* of points of the target space onto the latent space, but more importantly, we also gain a natural pull-back of the tangents in 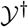 to tangents in 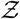. That is, given an element 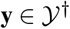 and a class *c* ∈ {1,…, *C*}, we define an average gradient of the pixels of *y* being predicted to class *c* by

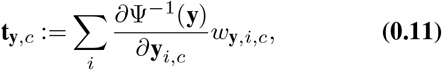

where *i* ∈ {1, …, *H* × *W*} is an indexing of pixels and *w*_**y**, *i*_ an importance weighting of the *i^th^* pixel in **y** to class *c*. If we denote 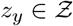 as the corresponding element satisfying Ψ(*z*_**y**_) = **y**, and 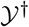 is a manifold, then for each pixel *i* and class *c*, the partial derivative 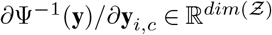 describes the sensitivity of *z*_**y**_ with respect to the *i^th^* pixel predicted as class *c*. Then, we take the weighted average over all pixels in the class of interest to construct an average sensitivity of *z_y_* with respect to class *c*, Eq. (0.11).

Using the *pull-back function,* Ψ^−1^, we define the a new latent space with untangled channels as follows:

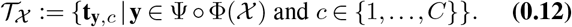

Let us consider a simple example to gain intuition into what the elements of the set above can represent. As from before, let *z*_**y**_ be the pull-back of a point **y** (see Fig. 2). Let Δ_*c*_(**y**) be a small perturbation of **y**, for example, a decrease in the prediction of class *c*. We pull-back Δ_*c*_(**y**) to *z*_Δ_*c*(**y**)__ using the pull-back Ψ^−1^. Then, the difference vector, *z*_*y*_ – *z*_Δ_*c*(**y**)__, is a coarse first order approximation of **t**_**y**, *c*_, that is, the vector pointing from *z*_Δ*c*(**y**)_ to *z*_**y**_. Such a directional vector can be constructed for each class *c*, (coloured arrows in Fig.2). It is not intuitive why a set of such tangent-like vectors, **t**_•_, should give a good latent space. Nevertheless, we will show through numerical experimental that 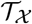 is a meaningful latent space, as it can be used to build saliency, predict functionals, like Intersection over Union (IoU), and more surprisingly, qualify unseen test data.

**Fig. 2.**
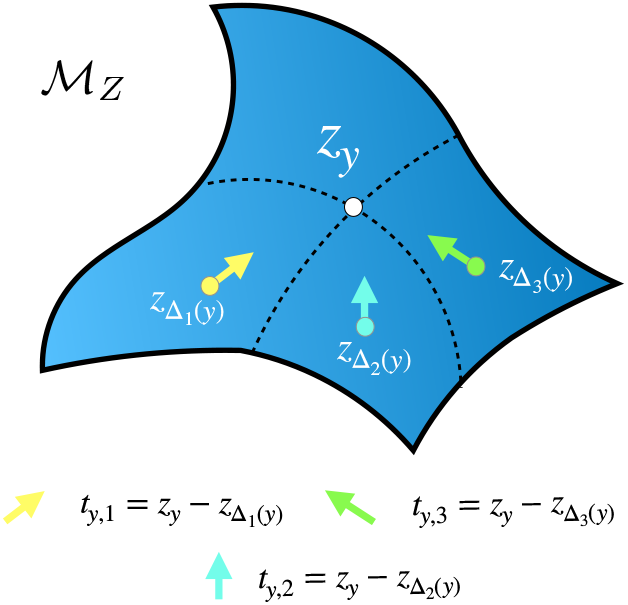
Cartoon. A sub-manifold 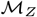 around the point *z_y_*. Three other points, coloured yellow, green and cyan, are pull-backs of *y* after perturbation with the function Δ_*c*_ to the class *c* ∈ {1, 2, 3}. The arrows are pull-back gradients for *y* and class c pointing to *z_y_*. Approximate direction to travel to reach *z_y_* from the perturbed state *x*_Δ_*c*^*y*^__.

## Methods

We performed numerical experiments to understand the Chimeric U-Net, which consists of a non-invertible encoder and an invertible decoder, in comparison to the standard (fully non-invertible) U-Net and the fully invertible U-Net. Furthermore, we study the latent space induced in the Chimeric U-Net and investigate its significance and impact. The following are the sub-modules used to construct the numerical experiments presented in the results.

### D. Architecture Models

We constructed three architectures: the standard U-Net (sU-Net), the invertible U-Net (iU-Net), and the Chimeric U-Net. We modified the architectures of the standard- and the invertible U-Net, such that, the encoder of the sU-Net and the decoder of the iU-Net matched with the encoder and decoder of the Chimeric U-Net, respectively. We present the detailed model representations in Appendix S-VII Fig. S-1.

The architecture of the sU-Net was based on the work by Ronneberger et al. (2). We modified their architecture in two ways: firstly, we expanded the input dimensionality to match the input channels with the output channels, which was done with a standard 2D convolution; secondly, we passed half the channels across the skip connection and passed the other half down to the successive layer, in contrast to (2), in which all channels were passed to both the skip connections and the layer below. Other than these changes, we used standard 2D convolutions with stride of two, kernel size of 3, and padding of one, to retain the spatial dimensions. Each convolution was followed by a Leaky ReLU with a negative slope of 0.01 and group normalisation (28).

The core of the iU-Net was based on the work by Etmann et al. (17). Two modification were made to the original architecture proposed, firstly we replaced the first and last non-invertible layer of the encoder and decoder in the architecture, with the `DWT` and `iDWT`, respectively, and secondly, to match the input and output channels, we inflated the input channels and set them to zeros prior to the DWT. Given the convolutions in the iU-Net only act on half the channels, we increased the number of additive coupling layers to four, to keep the total number of parameters in the architectures comparable.

The Chimeric U-Net is the combination of the encoder of the modified sU-Net and the decoder of the modified iU-Net. Once again, to match the overall number of parameters between the architectures, we reduced the number of additive coupling layers in the decoder units to be two. Descriptions of the models are given in Appendix S-VII.

All the U-Nets were build to have a depth of four. The architectures were implemented in `PyTorch` v1.9 (29). We used the *Cross Entropy Loss* function from the package as the loss function, furthermore, we used the Adam optimiser and trained with a learning rate of 10^-4^. We chose the model with the smallest loss on the respective validation set. The training was performed on a Nvidia Tesla P100-SXM2 and the subsequent analyses on an Intel Xeon Gold 6246 CPU.

### E. Datasets

We evaluate the U-Nets on two publicly available *Magnetic Resonance Imaging* (MRI) datasets: The Multimodal Brain Tumor Segmentation Challenge 2017 (BraTS), and the Zuse Insitute Berlin’s curated Osteoartheritis Initiative dataset 2018 (OAI ZIB).

#### BraTS (30), (31), (32)

The dataset contains densely annotated *low grade gliomas* (LGG) and *high grade gliobastomas* (HGG). For each patient: a T1 weighted, a post-contrast T1-weighted, a T2-weighted, and a *FLAIR* channel was available. Unlike other *BraTS* challenges, the 2017 edition contains only four target classes, namely: *background*, *edema* (ED), *non-enhancing tumor* (NET), and *enhancing tumor* (ET). The performance analysis was done for all classes, and also performed the analysis on the post-hoc merged class *tumor core*, which is the combination of ET and NET. The segmentation task was performed on 2D slices, consequently, 2D slices (240 × 240 pxls) of the 3D volumes were extracted. This was done by sampling three different slices from the inner 30% of each MRI scan. We chose a batch size of eight and chose to train for a maximum of 350 epochs. The data was split into five folds after reserving 10% of the data for testing. The splits were performed, such that samples of the same patient were not distributed among different data splits. In regards to data augmentation, flipping, blurring, random brightness, and elastic deformation were applied to the inputs during training (see Appendix S-I). We chose this particular dataset from the collection of *BraTS* datasets, because the chimeric- and invertible architectures did not require further channel adjustment to make the input channels match the number of target classes.

#### OAI ZIB (33), (34)

The dataset consists of roughly 500 3D MRI scans of human knees. The segmentation task was to predict five target classes, namely: *background* (BG), *femoral bone* (FB), *tibial bone* (TB), *femoral cartilage* (FC), and *tibial cartilage* (TC). As done for the *BraTS* dataset, 2D slices (384×384 pxls) were sampled from the inner 30% of each patients MRI. We chose a batch size of eight and chose to train for a maximum of 125 epochs. As for the BraTS dataset, the data was split into five folds after reserving 10% of the data for testing. During training the inputs were augmented by horizontal flipping, blurring and brightness changes, as mentioned above.

#### Tesselation dataset (35)

For the analysis of saliency maps we utilized the synthetic *Tesselation dataset* from the *Gap-Filling* problem. We trained a Chimeric U-Net for a maximum of 150 epochs with a batch size of eight. Thousand data samples were generated, each of size 240×240 pxls, and split into a train (80%), validation (10%) and test set (10%). In contrast to the Chimeric U-Nets trained for the *OAI ZIB* and *BraTS* dataset, no data augmentation was applied. The other hyperparameters were set as stated above (Methods D).

### F. Evaluation Metrics

We employed standard computer vision metrics for medical image segmentation, such as classwise and mean: *Intersection over Union* (IoU), *Recall*, and *Precision*.

### G. Pull-Back Gradient Approximations

Constructing the latent space 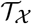 for a dataset is computationally challenging. For this reason, we chose to approximate the elements in 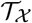 with coarse- and fine resolution approximations. In these computations our set 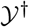 are the activation prior to applying the `softmax` (inline with Remark 0.1).

#### Coarse approximation

for 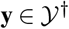 and class *c* ∈ {1,…, *C*}, our approximation of 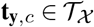 is given by,

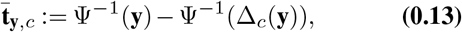

where we set Δ_*c*_ to be the operator that decreased the model prediction of class c by 70%. We denote the set of coarse approximation of the set 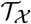 by 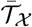. This approximation was used to generate the latent space embedding in the numerical experiments.

#### Fine approximation

We used a fourth-order numerical derivative approximation to estimate the partial derivative,

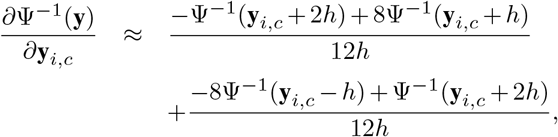

with *h* = 0.3 as the step size. We substituted the approximation of the partial derivative into Eq. (0.11) and use the predicted class probability as the weight *w*_**y**, *i, c*_. We denote the set of fine approximations of the elements of 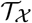 by 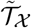.

### H. Vectorizing and Embedding the Pull-Back Gradients

Performing the analysis on the pull-back gradients, 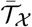, was not numerically feasible, and furthermore, we had to remove the spatial information, which was still encoded in them. To achieve this, we constructed a collapsing function,

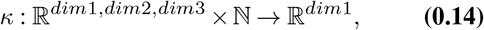

which takes an integer as input to determine how *dim*2 and *dim*3 are collapsed. The operation of this function is more suited to be presented as pseudo-code, hence, we present it as such in Appendix Algorithm 2. In this work, we chose the integer for collapsing to be the number of pixels which were positively predicted in the target. That is, let 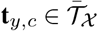, we performed a pull-back of *y* w.r.t. to class *c*. We define *k_y,c_* to be the number of pixels which were predicted (relative highest probability) to be class *c* in the target *y*. Then the collapse of **t**_*y, c*_ was given by,

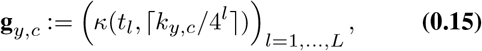

where *t_l_* is the *l^th^* element of **t**_*y,c*_. We refer to **g**_*y,c*_ as the *vectorized activation* of **t**_*y,c*_. This was performed for all element of 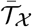 and we chose to visualize only the vectorized activation from only the deepest layer.

We used the t-SNE implementation in Python v3.7 (36) to visualize the vectorized activations. We chose *dim* to be 2, *perplexity* to be 150, *early exaggeration* to be 10, and the *correlation metric*. We normalised our data according to unit norm using norm *ℓ*_2_ before performing the embedding. The embedding was constructed after mixing the gradient pullbacks of 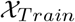 and 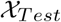.

### I. KNN Classifier

We used a k-nearest neighbours (kNN) implementation given in `scikit-learn` (37). We set *k* = 10 and constructed a function to map from the vectorized activations of 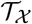 (as defined in Eq. (0.15)) to the IoU scores. The IoU score for an instance **g**_**y**, *c*_ is constructed by summing the IoU scores of the *k* nearest neighbours with the same class **g**_•, *c*_ and dividing by *k*. These calculations were performed layer-wise, and then for the final score, the average score over all layers were taken for robustness.

### J. Saliency Maps

We used the fine gradient approximations (Methods Eq. (0.14)) to study saliency maps. In order to obtain the saliency map, the Hadamard product of the elements from the gradient pull-backs, 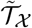, and pull-backs, 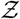, were summed, as in HiResCAM (19). The saliency maps were then sequentially upsampled, the corresponding pseudo-code is given in Appendix Algorithm 2.

## Results

In this section, we evaluate the qualitative and quantitative performance of the Chimeric U-Net compared to the iU-Net and the sU-Net for the BraTS- and OAI ZIB datasets. Furthermore, we present the approximations of the gradient pull-backs of the Chimeric U-Net for the BraTS- and the Gap Filling datasets, in the context of explainability.

### K. Combining a non-invertible encoder and an invertible decoder improves performance in comparison to the fully invertible U-Net

We evaluated the performance of the three architectures on BraTS 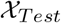 with five replicates and found that, the non-invertible sU-Net performed on average better with respect to the IoU and Precision scores over all the target classes (see Table 1). However, we also found an overlap of the 95% confidence intervals of the scores between the three architectures, suggesting that the performances were not necessarily significantly different. When we compared the Precision and Recall scores of the architectures for the classes NET and ET, we found that the iU-Net performed the worst. The observed decrease in Precision could be attributed to the lack of reduction of information by the iU-Net’s invertible encoder. Overall, we found that the performance scores of the Chimeric U-Net were closer to that of the sU-Net than the iU-Net, even though the number of trainable parameters for the Chimeric U-Net were closer to the iU-Net (≈ −10%) than the sU-Net (≈ +30%).

**Table 1.**
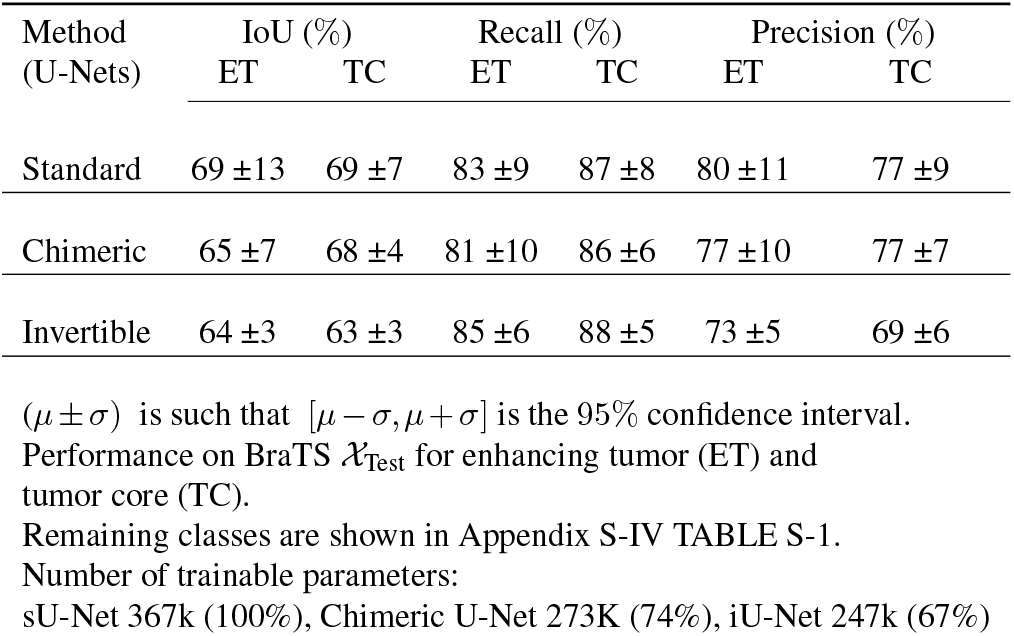

When we looked further into the shape of the individual predicted segmentations, we again found that the Chimeric U-Net prediction performance was between the sU-Net and the iU-Net. For example, the predictions of the sU-Net showed smoother contours along ED and TC (see segmentations in Fig. 3 (Row 2)). In contrast, the Chimeric U-Net and the iU-Net predictions showed rough boundaries (see segmentations in Fig. 3 (Row 3 and 4)). However, the iU-Net also produced artifacts (false positive predictions) on arbitrary positions on the image (see Fig. 3(•)) and rougher boundaries than the Chimeric U-Net (see segmentations in Fig. 3 (Column 4, 5 and 7)). Given the Chimeric U-Net and iU-Net have a similar decoder, we expected a closer behaviour, however, we found having a non-invertible encoder in the Chimeric U-Net, aided in reduction of the false positives rate and rough boundaries as observed in the iU-Net. We observed that all U-Nets struggled with detecting ED for low intensity samples (see yellow seg. in Fig. 3 (Column 6)). However, we found the predictions of the Chimeric U-Net to be closer to the sU-Net than the iU-Net (see yellow seg. in Fig. 3 (Column 2-5)). The Chimeric U-Net showed the precisest prediction for ED, whereas the sU-Net and iU-Net over-predicted it (see Fig. 3 (Column 2 and 5)), sometimes at regions where no ED was present (see yellow seg. in Fig.3 (‡)), suggesting that the Chimeric U-Net is much more robust against intensity fluctuations within its input.

**Fig. 3.**
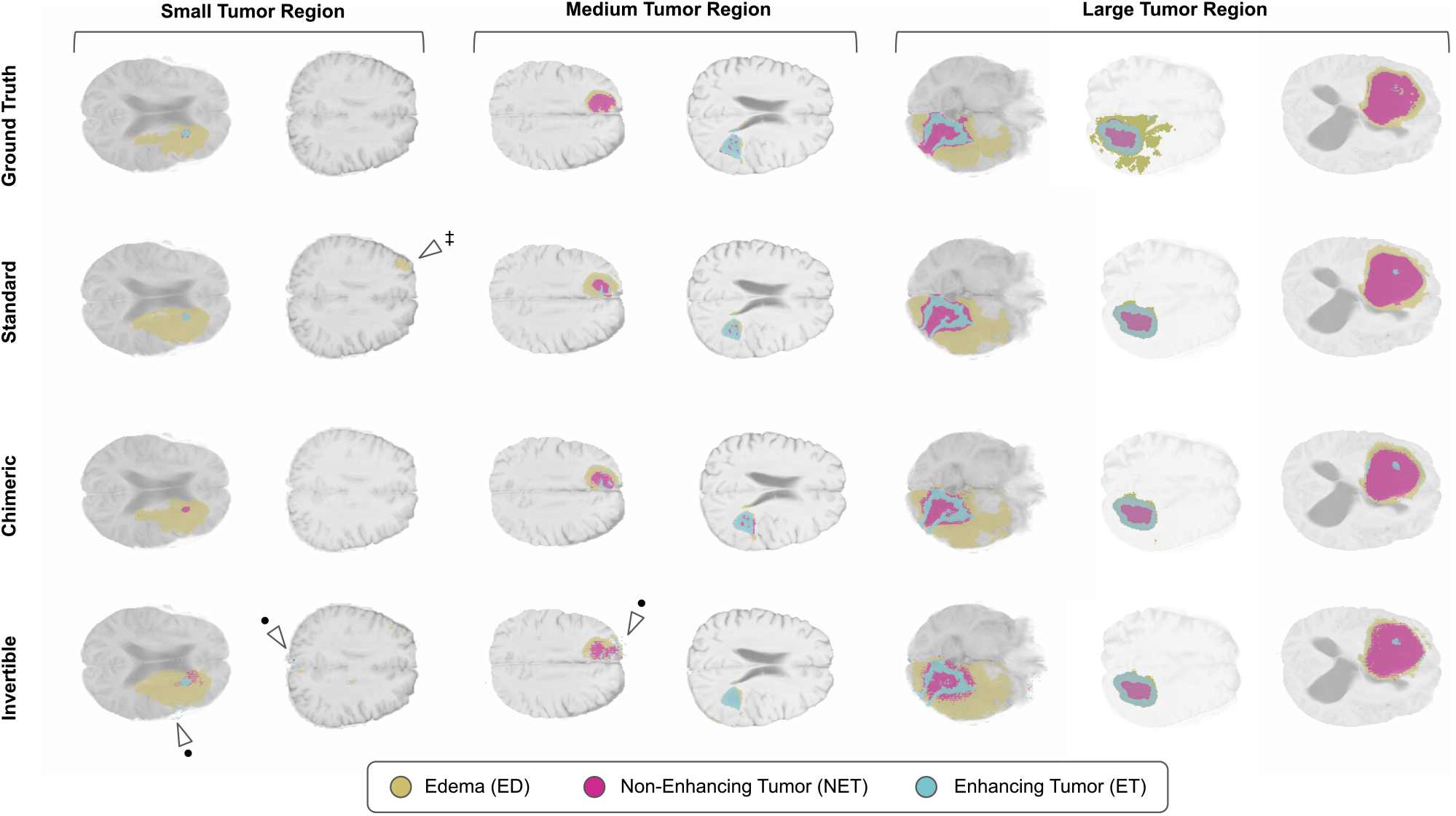
Segmentation results for samples from BraTS 2017 test dataset over input images (grey scale background) for the classes *edema* (yellow), *enhancing tumor* (blue) and *non-enhancing tumor* (purple). Rows show the ground truth and the models segmentation for standard, chimeric and non-invertible U-Nets. The columns show different samples further grouped by the severity of the tumor. The markers highlight observations discussed in the Results K.

In summary, having a non-invertible encoder aids in performing a better segmentation task and by this the Chimeric U-Net is at least as good as the iU-Net.

As a sanity check, we also trained the three architectures on the OAI ZIB dataset. Here we observed a similar trend in performance between the three architectures as with the BraTS dataset (see Table 2). That is, the sU-Net scores on average were better than that of the Chimeric U-Net and the iU-Net. In particular, we observed that the 95%confidence intervals of the Recall and Precision scores of the three architectures in the FC and TC classes were overlapping. Interestingly, the sU-Net had significantly better IoU scores ([74%, 76%]) in the FC class to the Chimeric U-Net ([71%, 73%]), but not to the iU-Net ([72%, 74%]), again showing that Chimeric U-Net is at least as good as the iU-Net. Looking closer at the individual predicted segmentations on the OAI ZIB test set, we observed the sU-Net to produce slightly smoother boundaries (best visible in Fig. 4 (Column 2)). Besides that, the overall differences between the architectures were subtle. The iU-Net was found to produce the best segmentations for FC, whereas the other architectures mostly over predicted it (see pink seg. in Fig.4 (⋆)). For TC, we observed it to be vice versa, i.e. the iU-Net over predicted TC. This can be seen for the second sample of Fig. 4, for which only the Chimeric U-Net and sU-Net predicted correctly the absence of TC (see green seg. in Fig. 4 (‡)). Furthermore the Chimeric U-Net often showed more precise predictions for TC (see green seg. in Fig. 4 (■)). Lastly, for the sU-Net we noticed sometimes false predictions in bright spots of the images (see orange seg. in Fig. 4(•)).

**Fig. 4.**
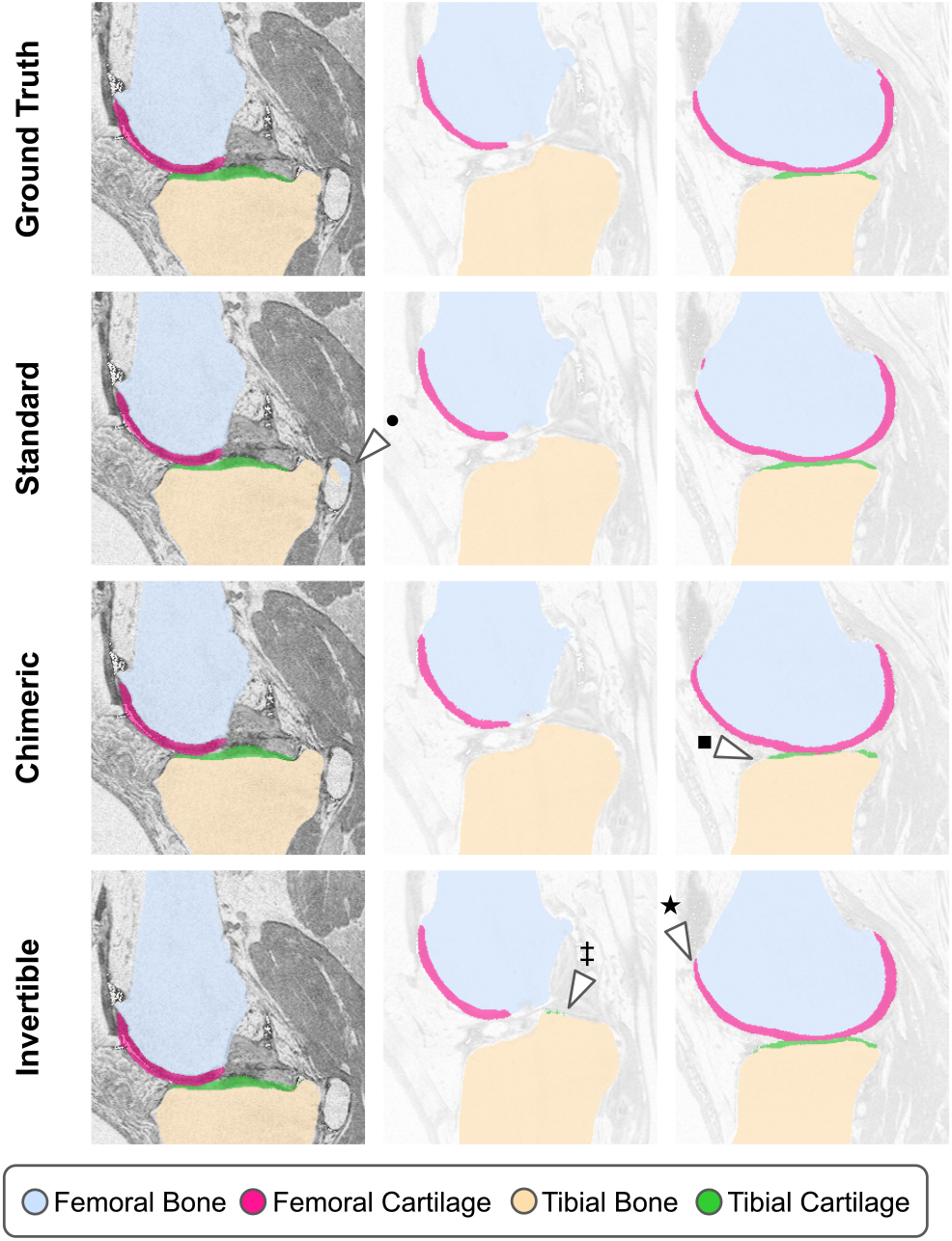
Segmentation results for samples from OAI ZIB test dataset over input images (grey scale background) for the classes *femoral bone* (blue), *tibial bone* (orange), *femoral cartilage* (pink) and *tibial cartilage* (green). Rows show the ground truth and the models segmentation for standard, chimeric and non-invertible U-Nets. The markers highlight observations discussed in the Results K.

**Table 2.**

| Method<br>(U-Nets) | IoU (%) |  | Recall (%) |  | Precision (%) |  |
| --- | --- | --- | --- | --- | --- | --- |
|  | FC | TC | FC | TC | FC | TC |
| Standard | 75 ± 1 | 61 ± 1 | 85 ± 3 | 74 ± 6 | 87 ± 3 | 78 ± 5 |
| Chimeric | 72 ± 1 | 58 ± 2 | 82 ± 3 | 70 ± 3 | 86 ± 1 | 78 ± 4 |
| Invertible | 73 ± 1 | 56 ± 3 | 84 ± 1 | 66 ± 5 | 84 ± 1 | 79 ± 3 |

( $\mu \pm \sigma$ ) is such that $[\mu - \sigma, \mu + \sigma]$ is the 95% confidence interval. Performance on OAI ZIB $\mathcal{X}_{\text{Test}}$ for femoral cartilage (FC) and tibial cartilage (TC).
Remaining classes are shown in Appendix S-IV TABLE S-2.
Number of trainable parameters:
sU-Net 574k (100%), Chimeric U-Net 427K (74%), iU-Net 386k (67%)

In summary, the Chimeric U-Net is on par –at times better– with the iU-Net for segmentation tasks over the BraTS and ZIB OAI datasets. The non-invertible encoder is needed to increase precision, through the removal of redundant information. Combining a non-invertible encoder with an invertible decoder, as in the Chimeric U-Net, gave results closer to the purely non-invertible setting. Nevertheless, we observed a minor performance drop of the Chimeric U-Net to the sU-Net, but the trade-off for this is in the explainable properties, which comes from the invertible decoder of the Chimeric U-Net, which we demonstrate next.

### L. The vectorized activations contain a meaningful structure w.r.t. the target classes

We used the coarse grained derivative approximation (Eq. (0.13)) to compute the pull-back gradients of the individual segmentations (see Methods G – H) of the BraTS dataset and then used the t-SNE embedding to visualize their corresponding vectorized activations (see Fig.5).

**Fig. 5.**
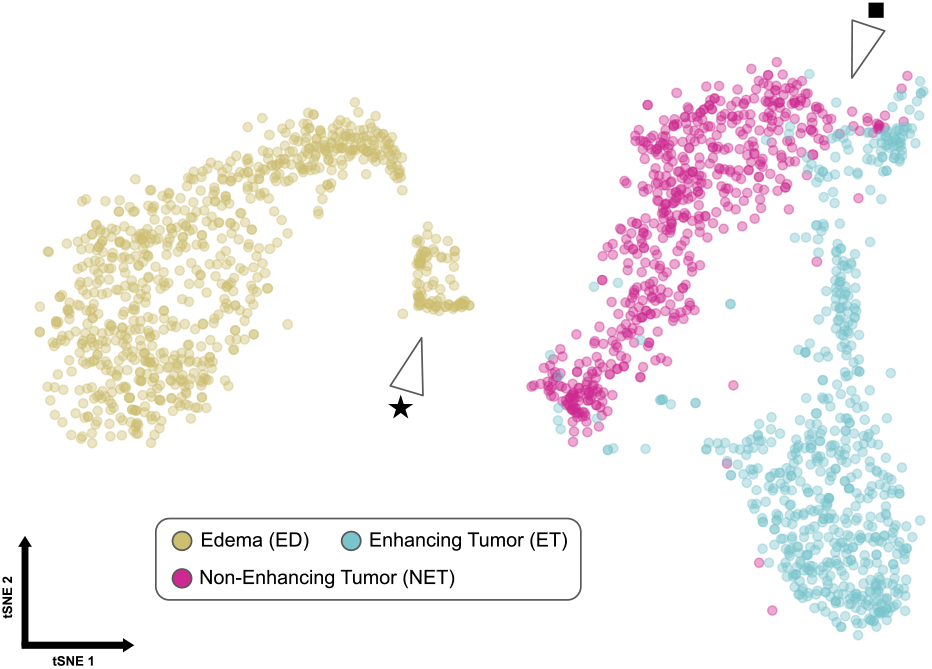
Visualization of t-SNE embedding of vectorized activations for BraTS 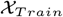 performed by Chimeric U-Net. Colors display target classes, namely *ED* (yellow), *ET* (blue) and *NET* (purple). Each point represents a single pull-back for one class of target sample. The markers highlight observations discussed in the Results M.

In the embedding of the train dataset 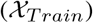, we observed that the three target classes: NET (purple), ET (blue), and ED (yellow), clustered distinctly from each other, with slight overlap of the two tumor types, i.e. NET and ET (see Fig. 5 (■)). Furthermore, we observed in the embedding, that the ED points had a small sub-cluster closer to the other classes (see Fig. 5 (⋆)), suggesting that segmentation of ED corresponding to these points was perhaps different to the remaining ED points.

To understand the meaning of the spatial distances between the vectorized activations, we overlayed the IoU scores of the respective class prediction on the t-SNE visualisation (see Fig.6). Surprisingly, we saw that the smaller ED cluster points and the overlapping NET and ET points showed low IoU scores (see Fig.6 (⋆)). Additionally, instances of images from the mixed regions of NET and ET were falsely predicted as the other class (see Fig.6 (‡)),as expected. Lastly, we saw that there were regions within each class cluster, where all the points in the spatial proximity had similar high scores (e.g. Fig.6 (•)). Recall, that we normalized the pull-back gradients by the size of the class positives (see Eq. (0.15) to reduce the influence of varying sizes of predictions.

**Fig. 6.**
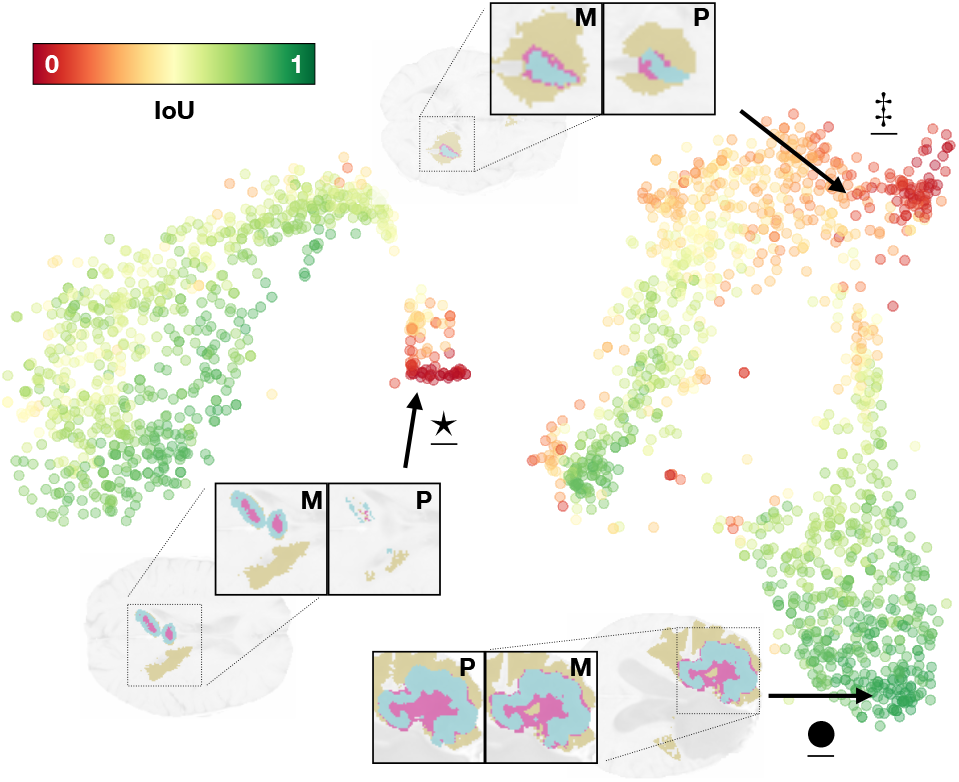
Visualization of t-SNE embedding of the vectorized activations for BraTS 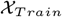 colored by evaluated per class IoU, performed by the Chimeric U-Net. The brain scans depict exemplary samples from different positions within the embedding, overlayed with their respective model’s prediction (**P**) and the ground truth mask (**M**). The markers highlight observations discussed in the Results M.

In summary, we found that the vectorized activations of the target class predictions cluster in a meaningful way, driven by the class membership and the prediction performance.

### M. The vectorized activations form a meaningful latent space to infer the accuracy of unseen data

We investigated if the IoU score relationship to the spatial proximity also translated to unseen data 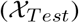. For this reason, we employed a simple kNN-classifier (see Methods I) to estimate the IoU score corresponding to a vectorized activations based on the local proximity to the vectorized activations from the training data. When overlayed onto the embedding of the vectorized activations of the test dataset the evaluated IoU score and the kNN estimated IoU scores, we found that they were visually similar (see Fig.7a–b). Furthermore, plotting the estimated verses the evaluated IoU scores, we observed that the estimated IoU scores were a weak upper bound to the evaluated scores, and that the two became positively correlated for higher score values (Fig.7 c).

**Fig. 7.**
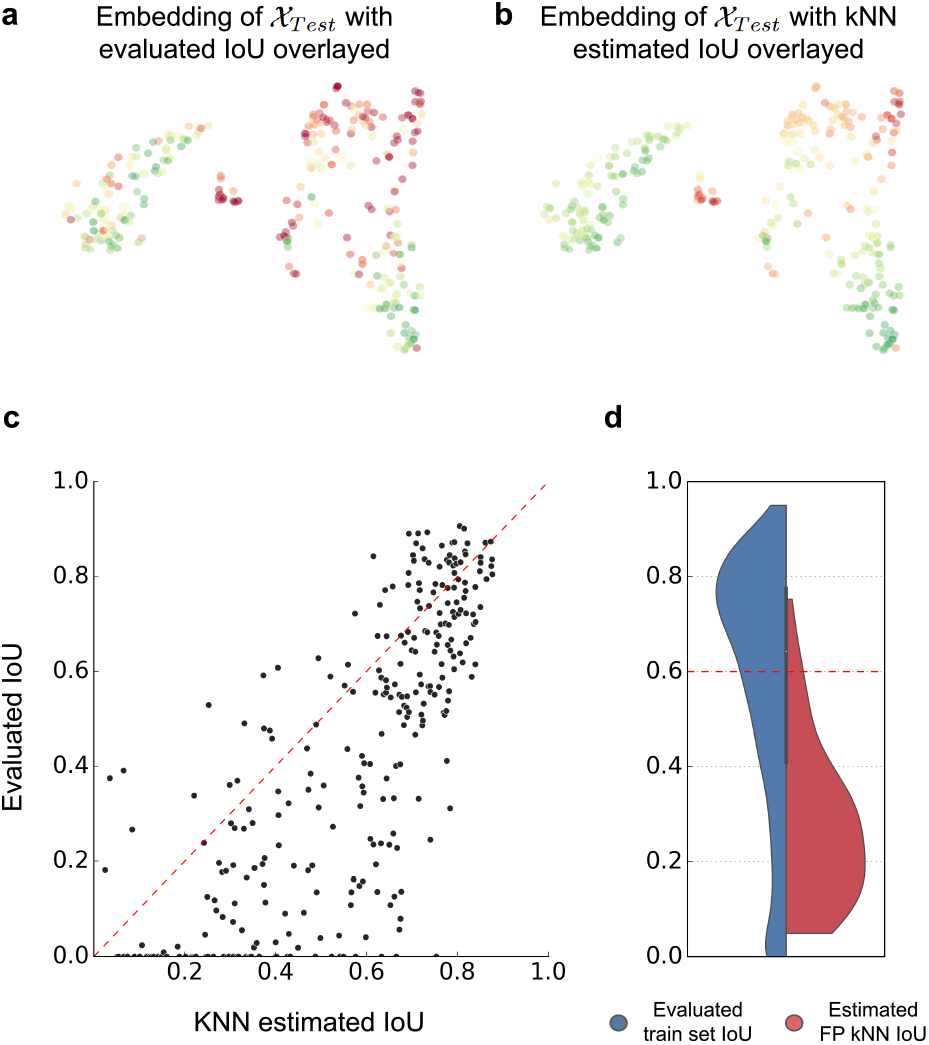
Top: Visualization of t-SNE embedding of vectorized activations for BraTS 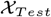 performed by the Chimeric U-Net. Colors display (a) the evaluated IoU scores (b) and the IoU scores estimated with the kNN-classifier. Bottom: Comparison of evaluated and estimated IoU. (c) Overall fit of kNN-classifier-estimate versus evaluated IoU for 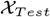 and (d) contrast of estimates of the kNN-classifier for false positive predictions to the evaluated IoU scores for 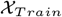.

Given the pull-backs were made for all classes predicted by the Chimeric U-Net, naturally, the *false positives* (FPs) were also pulled back. Hence, we observed 58 vectorized activations which had positive estimated IoU, but scored a value of zero, under this metric (Fig.7 c (x-axis)). Even though these were FPs, the kNN estimate gave only roughly 5%of the vectorized activations a score of above 60%. (see Fig.7d). In addition, the majority of the FPs had a low estimated score of roughly 20%. When considering the evaluated IoU scores for 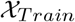, it can be seen clearly that those low scores correctly represented poor prediction performance.

In summary, the clustering of the vectorized activations of the training dataset encodes how confident the architecture is in its class predictions. More importantly, the vectorized activations of the training data can be used to give an estimate on the true accuracy of the prediction of unseen data.

### N. Pull-back gradients from the invertible decoder can be used to compute saliency maps of segmentations

Saliency maps are the state-of-the-art approach for explainability in classification tasks. We investigated if the decoder in the Chimeric U-Net could be used to also generate saliency maps of the pixel predictions in the segmentation task. We found that both, the BraTS and OAI ZIB, dataset sets produced trivial saliency maps, suggesting that the segmentation task was driven by texture rather than by context (results not shown). Hence, to detect context decisions made by the Chimeric U-Net, we utilized the synthetic *tessellation dataset* from the *Gap-Filling* problem. In this task, the architecture had to segment lines in the input image, but additionally also had to fill the gaps between incomplete lines, for which the gap pixels are indistinguishable from background pixels (Methods D).

The Chimeric U-Net performed well on the *Gap-Filling* problem, as lines were correctly segmented with an IoU score of 86%. We choose a representative source image from the test set to be segmented. On this image, we picked three points of interest: a background pixel, a line pixel, and a gap pixel (see Fig.8 a (dashed boxes)). For these pixels, we computed their respective saliency maps, that is, highlighting the regions in the source image which aided in these pixels to be classified as a line pixel (Methods I).

**Fig. 8.**
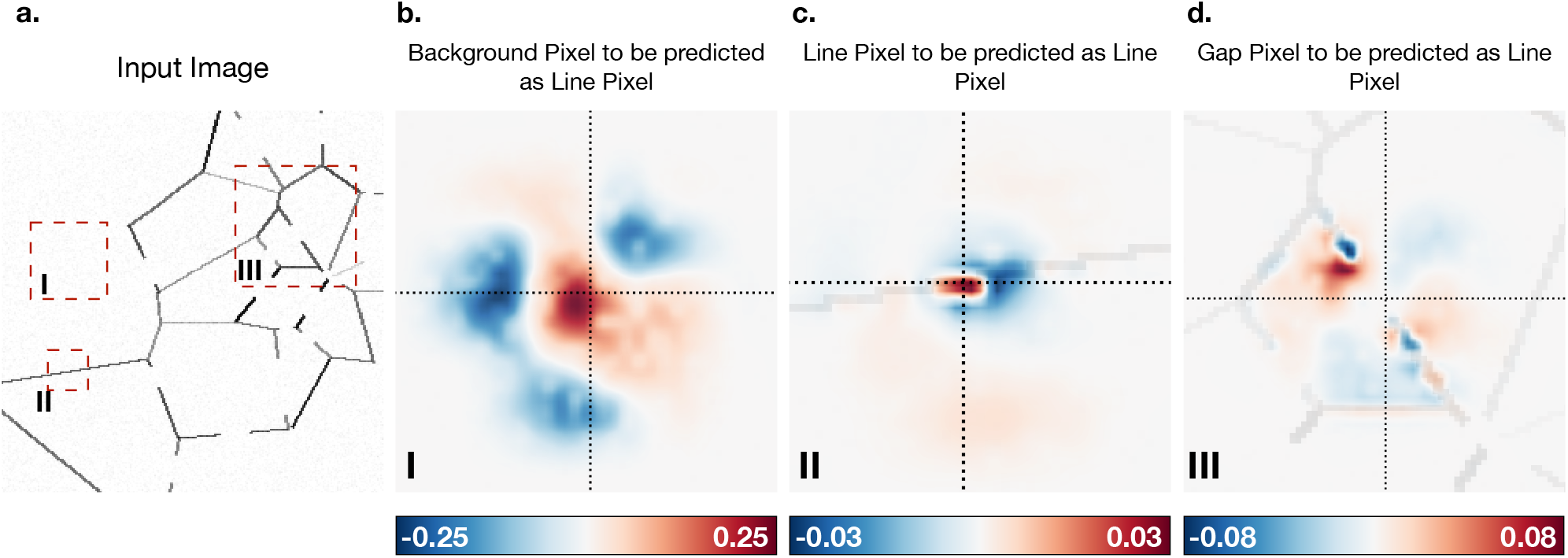
Saliency maps for different pixels (indicated by dashed cross) of sample from gap dataset. Only for pixels that had to be filled in order to fill a gap relied only on context information. The coloring shows the positive (blue) and negative (red) shaping the saliency maps.

Firstly, for the background pixel, we saw that the highest saliency was in proximity of the pixel itself and the signal diffused outwards, mimicking the receptive field (see Fig.8 b). Secondly, for the the line pixel, we saw a strong saliency signal on the line segment, and some weaker signal radiating away from the line (see Fig.8 b). Lastly, for the gap pixel, we saw that the saliency was highest at the ends of the lines at either end of the gap, with nearly no signal at the pixel, showing that the prediction is explained by the context of the edges on either side of the of the gap.

In summary, the pull-back gradients from the invertible decoder can be easily used to construct saliency maps.

## Discussion

We proposed the Chimeric U-Net, a novel explainable CNN architecture based on the standard U-Net, but with an invertible decoder that could successfully solve segmentation tasks with built-in explainability. The discussion points below are structured to match the points presented in the results, that is, we discuss the impact and limitations of: the technical aspects of the architecture, the class disentangled latent space embedding, and lastly, the pull-back gradients based saliency maps.

We showed that the Chimeric U-Net performed en part with the purely invertible U-Net on two biomedical image segmentation tasks, suggesting that the non-invertible encoder has little negative impact on the performance. In contrast, the Chimeric U-Net performed slightly worse compared to the sU-Net, suggesting that the invertible decoder does have some negative effect on performance. Here, we proposed a minimal architecture to demonstrate the concept of using an invertible decoder, since technical and practical aspects of architecture analysis were not direct focus of this work. We speculate that the decrease in performance shown in this work is primarily due to the inverse discrete wavelet transform (`iDWT`) that we introduced to obtain the total invertibility of the iU-Net. Given the `iDWT` has no trainable parameters, this enforces the Chimeric U-Net to learn the segmentation in the steps prior to its application. We performed post hoc experiments, in which the `iDWT` was exchanged with non-invertible standard convolutions, and observed that this enhanced the segmentation quality, at the cost of the model’s explainability (results not shown). Naturally, higher order basis functions for the `iDWT` can be chosen, and also, it can be exchanged with other invertible neural functions. In practice, this should be tailored to the dataset of interest. Given the flexibility of the encoder in the Chimeric U-Net, the optimal choice of the architecture’s encoder and its influence on the information bottleneck over the skip connections is still to be explored. Another technical advantage we did not exploit in this work is the memory consumption savings given by the invertible components as presented by Etmann et al. This would be a natural future direction of research to construct bigger Chimeric U-Net architectures.

Constructing an embedding that can estimate the quality of the data for which no ground truth is available has tremendous impact in fields like healthcare, in which large amounts of data are generated (38). The use of the derivative of the pullbacks with respect to the classes (pull-back gradients) was a fruitful idea for constructing a class-disentangled latent space embedding. We could see that in the skip connections of the Chimeric U-Net architecture, different classes were passed over different activations, and over the same activations when there was a confusion in delineating the classes. This feature splitting was anticipated given the bottleneck caused by the non-invertible decoder, and since clustering of the latent space with respect to the classes was already previously observed in classical classification tasks. However, what came as a surprise was how the distribution of points inside the clusters were correlated with the IoU, indicating that there is a way to estimate the performance of unseen data through the latent space. This could naturally be used to check for generalisability of the architecture. For example, if the architecture was trained on a particular genealogical demographic and then was applied to another demographic, then we could check where the prediction of the two groups are located in the latent space and interpret hidden biases. A line of questioning which we didn’t consider in this work was to investigate if the pull-back gradients could detect subclasses within a poorly labelled class. For example, if we had “enhancing tumor” and “non-enhancing tumor” labelled as a meta class “tumor”, then we would want the pull-back gradients of the class “tumor” to be split into at least two clusters, where one cluster represented patients who had more enhancing tumor over nonenhancing, and vice versa. Another question we only explored in part was how the correlations between the classes form over the training period. By using a simple correlation measure, we can investigate which classes are correlated within the Chimeric U-Net architecture. This knowledge can be used for better training or model debugging, for example, by increasing specificity of the class labels, assessing data quality and normalization, and monitoring the training progress of the network (39), (40). Hence, studying the pull-back gradients and their effectiveness to explain the overall behaviour of the architecture is a natural future research direction.

Historically, saliency maps were highlighted regions on an image where a viewer was focusing in visual experiments. This concept was translated into modern XAI, and since has been a major tool and topic of much ongoing debate. The latter stems from the lack of ground truth and quantitative metrics, thus making correctness and faithfulness of saliency methods contentious (24), (25), (41), (26), (27), (23). However, saliency maps are important for human interpretation, hence, we wanted to investigate if the question (XAI *β*) could also produce a saliency map. To circumvent the criticism around validation of the saliency maps, we chose to use the gap-filling problem, where a mathematical proof was already given on where the saliency of a classification of a line segment should lay in the source image. We could visually verify that the saliency maps made from (XAI *β*) did highlight the expected regions in the input image, suggesting that (XAI *β*) could also construct meaningful contextual saliency maps. The saliency maps in this work were constructed for only the positive class predictions, however, a future direction would also be to consider the pull-back gradients of the false negative predictions and investigate their saliency maps. Lastly, recalling from the introduction, we do not give a comparison between (XAI *α*) and (XAI *β*) in this work. With the emergence of metrics and ground truth for saliency maps, a clear future direction would be to investigate and categorize the various saliency maps arising from XAI questions.

In conclusion, the Chimeric U-Net arises from the principle of enforcing explainability into the architecture, rather than studying explainability of a general architecture. Through the invertibility of the decoder we could inherently produce both global- and local explainability through class embeddings and saliency maps, respectively. The notion of having an invertible decoder can also be translated to other Deep Learning tasks, where explainability is currently scarce. We believe that the Chimeric U-Net architecture is a step in the right direction towards growing confidence, reliability, and trust in Deep Learning approaches in healthcare.

## Notes

### Competing Interest Statement

The authors have declared no competing interest.

### Summary of Updates

Added Code Link

