## Supplemental Material for "Chimeric U-Net – Modifying the standard U-Net towards Explainability" (from another version of this preprint: version 1, 2022-12-02)

### Supplementary Note 1: Data Augmentation

For the data augmentation routine of the the BraTS- and OAI ZIB dataset we utilized *Albumentations* (1). For simplicity, we denote the probability of applying a transformation during training to a sample by $p$. For the BraTS dataset we performed vertical and horizontal flips ($p = 0.5$), blurring ($p = 0.5$, $blur\_limit = 7$), random brightness ($p = 0.3$), and elastic deformations ($p = 0.3$, $\alpha = 120$, $\sigma = 6$, $\alpha\_affine = 3.6$). For the OAI ZIB dataset we performed only horizontal flips ($p = 0.5$), blurring ($p = 0.5$, $blur\_limit = 7$), and random brightness ($p = 0.3$).

### Supplementary Note 2: Predict IoU by Object Size

In this section we dive deeper into the kNN-scoring presented in the main text, by first, comparing the evaluated- and kNN-estimated IoU scores for the BraTS training set, secondly, comparing the kNN-estimate to a simple IoU estimate by the size of the predicted objects, and lastly, comparing the evaluated and kNN-estimated IoU scores for the the OAI ZIB testing set.

We observed the kNN-estimate to show a tight fit for the BraTS training set, when compared to the corresponding evaluated IoU scores (see Fig.1).

Naturally, larger objects are easier to predict, as more information can be encoded within their surface, in contrast to objects with the size of only a few pixels. Consequently, we investigated if the object size is a good predictor for the evaluated IoU and compare it to the proposed kNN-estimate. Non surprisingly, we found a positive correlation between object size and evaluated IoU of the training set (see Fig. 2) and testing set (see Fig. 3).

However, we found, the size-estimate to be a bad predictor, especially when compared to the kNN-estimate, as the evaluated IoU scores vary strongly for either small or large objects. To quantify this, we computed the Pearson correlation coefficient (PCC) with respect to the evaluated IoU scores, for both the kNN-estimate and the size-estimate, for which the former showed a roughly 15% higher score. The results strengthen the finding, that the extracted latent space encodes information beyond simple pixel magnitudes. We tested the kNN-classifier also on the OAI ZIB dataset, aiming to estimate the IoU of the testing set. We found the results to be a very good estimate for the evaluated IoU scores (see Fig. 4).

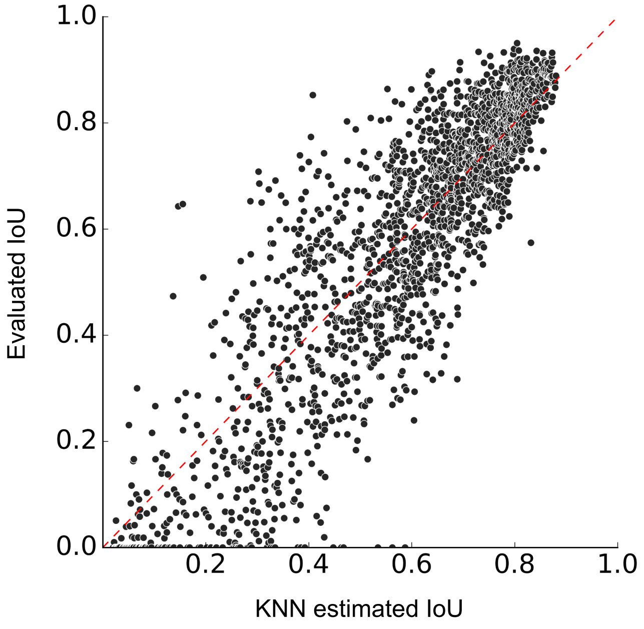


**Fig. 1.** Comparison of evaluated IoU scores versus its kNN-estimates on the BraTS training set.

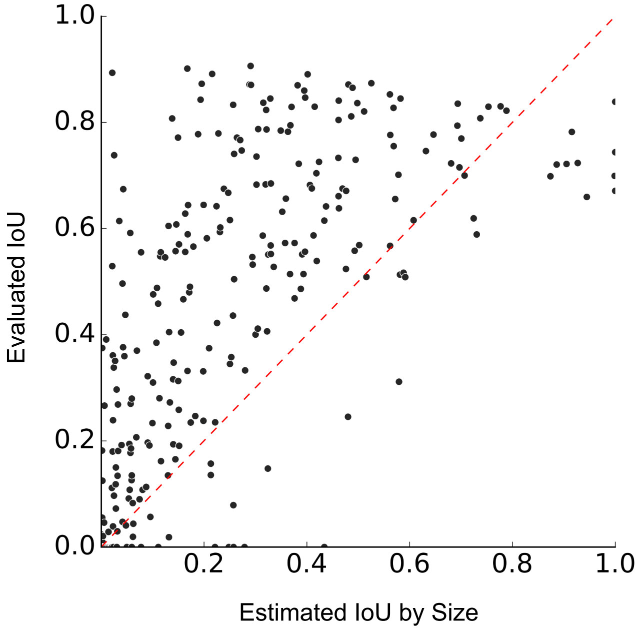


**Fig. 2.** Comparison of evaluated IoU scores versus its size-estimates on the BraTS training set.

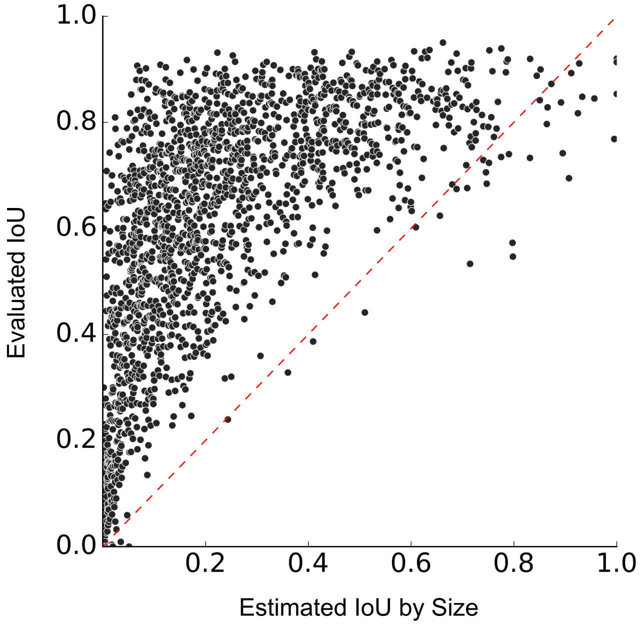


**Fig. 3.** Comparison of evaluated IoU scores versus its size-estimates on the BraTS testing set with an PCC of 0.654 (PCC of 0.789 for kNN-estimate).

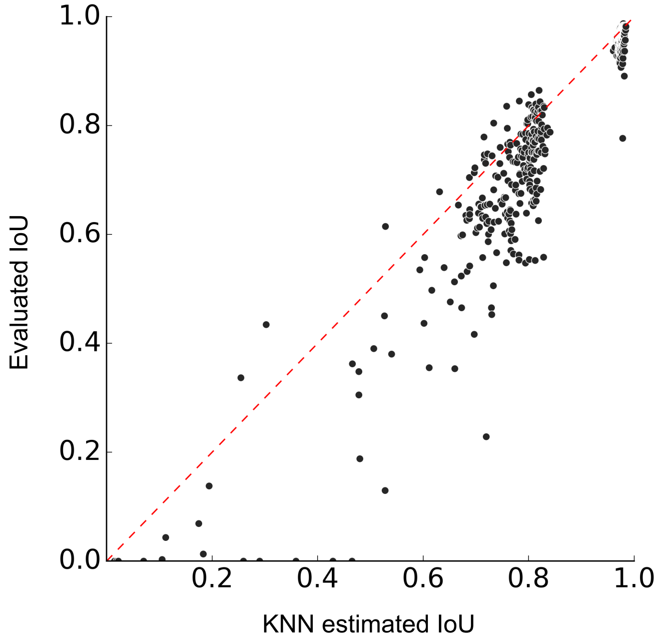


**Fig. 4.** Overall fit of the kNN-classifier estimate versus evaluated IoU for the OAI ZIB $\mathcal{X}_{Test}$ of the Chimeric U-Net.

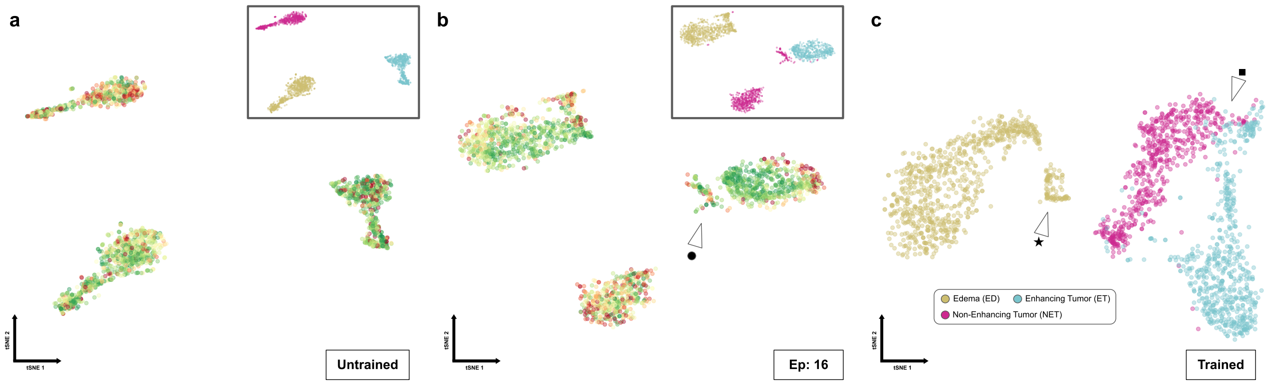


**Fig. 5.** T-SNE trajectory for different training steps of the Chimeric U-Net for the BraTS training set.

### Supplementary Note 3: Monitoring the training progress through the embedding

By creating t-SNE trajectories for different time steps of the Chimeric U-Net we monitored the training process for the BraTS training set (see Fig. 5). We observed no inter-class correlation in the embedding space for the untrained Chimeric U-Net. Furthermore, points, predicted poorly by the fully trained model, were spread disordered within the class cluster boundaries of the embedding. After 16 epochs, we found that a fraction of the embedded NET cluster moved closer towards the ET cluster (see Fig. 5 (●)). For the moved fraction, the trained Chimeric U-Net showed high prediction performance, while the larger part of the NET cluster, which kept being separated, showed no structure w.r.t. the IoU score.

### Supplementary Note 4: Pseudocodes

In this section we show the corresponding pseudocodes, named in the main paper.

**Algorithm 1:** Vectorize Activations

**Input:** Trained Chimeric U-Net : M
**Output:** **g**
**Data:** $\mathcal{X}$

/* iterate over all training samples */
$i \leftarrow 0$
**for** $i$ *in* $|\mathcal{X}|$ **do**
  $y \leftarrow M(\mathcal{X}[i])$
  $c \leftarrow 0$
  **for** $c$ *in* $|C|$ **do**
    $n\_positives \leftarrow \sum((\arg\max_c(y)) == c)$
    /* $z_{\mathbf{y}} - z_{\Delta_c(\mathbf{y})}$ */
    $\bar{\mathbf{t}}_{\mathbf{y},c} \leftarrow \text{GradientApprox}(M, y, c, \Delta_c)$
    $\mathbf{g}_{\mathbf{y},c} \leftarrow \kappa(\bar{\mathbf{t}}_{\mathbf{y},c}, n\_positives)$
    $c \leftarrow c+1$
  $i \leftarrow i+1$

**Algorithm 2:** Collapsing Function $\kappa$

**Input:** $\bar{\mathbf{t}}_{\mathbf{y},c}$, n_positives
**Output:** $\mathbf{g}_{\mathbf{y},c}$

/* iterate over all layers */
$l \leftarrow 0$
**for** $l$ *in* $|\bar{\mathbf{t}}_{\mathbf{y},c}|$ **do**
  $k \leftarrow \lceil n\_positives/4^l \rceil$
  /* filter k highest elements per channel */
  /* $CxNxM$ -> $CxUxV$, where $u \leq n, v \leq m$ */
  $*\bar{\mathbf{t}}_{\mathbf{y},c} \leftarrow \text{FILTER}(\mathbf{t}_{\mathbf{y},c,l,k})$
  /* $CxUxV$ -> $C$ */
  $\mathbf{g}_{\mathbf{y},c} \leftarrow \frac{1}{Z}\sum_u \sum_v *\mathbf{t}_{\mathbf{y},c}$, where $Z = U * V$

**Algorithm 3:** Sequential Upsampling

**Input:** $Features : \mathcal{Z}_l, Gradients : \tilde{\mathcal{T}}_l$
**Output:** Saliency Map S

$l \leftarrow 1$
**for** $i$ *in* $|\mathcal{Z}_l|$ **do**
  /* Double the spatial dimensions */
  $\mathcal{T}_l \leftarrow Rescale(\tilde{\mathcal{T}}_l, 2)$
  $\mathcal{T}_l \leftarrow \mathcal{T}_l \odot \mathcal{Z}_{l-i}$
  $l \leftarrow l+1$
$S \leftarrow \mathcal{T}_l$

**Table 1**

| Method (U-Nets) | IoU (%) ED | IoU (%) NET | Recall (%) ED | Recall (%) NET | Precision (%) ED | Precision (%) NET |
|---|---|---|---|---|---|---|
| 2Standard | 57 ±7 | 44 ±7 | 77 ±10 | 68 ±10 | 69 ±9 | 56 ±10 |
| Chimeric | 58 ±4 | 41 ±3 | 77 ±5 | 64 ±9 | 72 ±3 | 54 ±6 |
| Invertible | 51 ±8 | 37 ±7 | 73 ±7 | 66 ±8 | 64 ±13 | 46 ±8 |

$(\mu \pm \sigma)$ is such that $[\mu - \sigma, \mu + \sigma]$ is the 95% confidence interval.
Performance on BraTS $\mathcal{X}_{\text{Test}}$ for edema (ED) and
non-enhancing tumor (NET).
Number of trainable parameters:
sU-Net 367k (100%), Chimeric U-Net 273K (74%), iU-Net 247k (67%)

**Table 2**

| Method (U-Nets) | IoU (%) FB | IoU (%) TB | Recall (%) FB | Recall (%) TB | Precision (%) FB | Precision (%) TB |
|---|---|---|---|---|---|---|
| Standard | 96 ±0 | 97 ±0 | 98 ±0 | 98 ±0 | 98 ±0 | 98 ±1 |
| Chimeric | 96 ±0 | 97 ±0 | 98 ±0 | 98 ±0 | 98 ±0 | 98 ±0 |
| Invertible | 96 ±0 | 96 ±0 | 98 ±1 | 99±0 | 98 ±1 | 98 ±0 |

$(\mu \pm \sigma)$ is such that $[\mu - \sigma, \mu + \sigma]$ is the 95% confidence interval.
Performance on OAI ZIB $\mathcal{X}_{\text{Test}}$ for *femoral bone* (FB) and
*tibial bone* (TB).
Number of trainable parameters:
sU-Net 574k (100%), Chimeric U-Net 427K (74%), iU-Net 386k (67%)

### Supplementary Note 5: Performance on omitted Classes

In this section, we report the summary statistics for the sU-Net, iU-Net, and Chimeric U-Net for the classes, omitted in the main paper.

### Supplementary Note 6: Embedding of Knee Dataset

We used coarse grained derivative approximation to compute pull-backs to extract a latent space for the OAI ZIB dataset, for which the Chimeric U-Net showed very good prediction performance. Within the latent space the classes were disentangled and showed, in contrast to the latent space extracted for BraTS dataset, no inter-class mixing. We further observed in the t-SNE embedding (see Fig. 6 (a)), that the clusters have arranged similar to the segmentation masks, with the cartilage classes being in between the two bone classes.

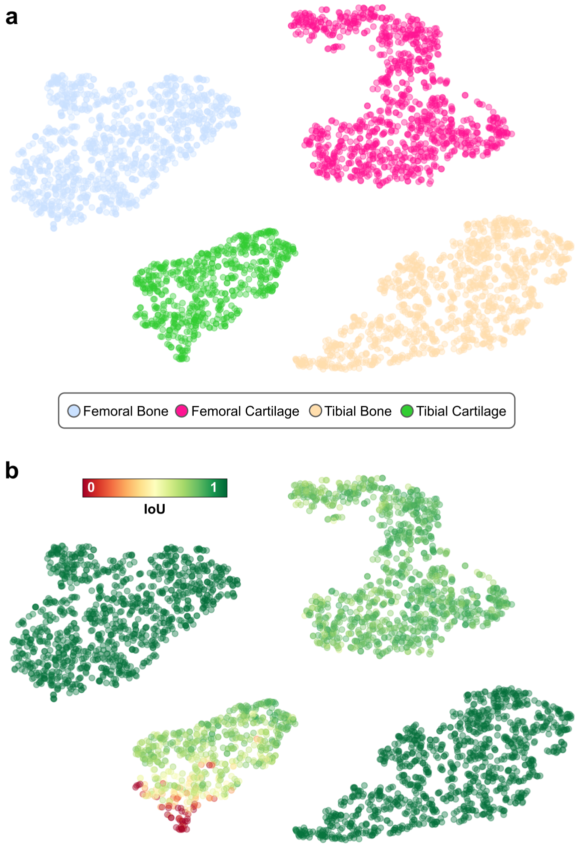


**Fig. 6.** Visualization of t-SNE embedding of vectorized activations for OAI ZIB $\mathcal{X}_{Train}$ performed by the Chimeric U-Net. Colors display (a) target classes and (b) IoU score for each pull-back.

When overlaying the evaluated IoU scores on top of the t-SNE embedding for the pull-backs (see Fig. 6 (b)), we found, as for the pull-backs of the BraTS dataset, the IoU score to be encoded within the latent space (best visible for the pull-backs of *tibial cartilage*). Lastly, we observed that during training the distances of the clusters shrank (results not shown), suggesting that the model learns to correlate the target classes in order to enhance the predictivity.

### Supplementary Note 7: Model Description

In this section we list the three model structures used for the analysis in the main paper.

| Modules | In x Out x H x W | Parameters |
|---|---|---|
| up.weight | [4, 16, 3, 3] | 576 |
| up.bias | [16] | 16 |
| encoder.layers.0.block.0.F2.conv.weight | [16, 16, 3, 3] | 2304 |
| encoder.layers.0.block.0.F2.norm.weight | [16] | 16 |
| encoder.layers.0.block.0.F2.norm.bias | [16] | 16 |
| encoder.layers.0.block.1.F2.conv.weight | [16, 16, 3, 3] | 2304 |
| encoder.layers.0.block.1.F2.norm.weight | [16] | 16 |
| encoder.layers.0.block.1.F2.norm.bias | [16] | 16 |
| encoder.layers.1.block.0.F2.conv.weight | [8, 32, 3, 3] | 2304 |
| encoder.layers.1.block.0.F2.norm.weight | [32] | 32 |
| encoder.layers.1.block.0.F2.norm.bias | [32] | 32 |
| encoder.layers.1.block.1.F2.conv.weight | [32, 32, 3, 3] | 9216 |
| encoder.layers.1.block.1.F2.norm.weight | [32] | 32 |
| encoder.layers.1.block.1.F2.norm.bias | [32] | 32 |
| encoder.layers.2.block.0.F2.conv.weight | [16, 64, 3, 3] | 9216 |
| encoder.layers.2.block.0.F2.norm.weight | [64] | 64 |
| encoder.layers.2.block.0.F2.norm.bias | [64] | 64 |
| encoder.layers.2.block.1.F2.conv.weight | [64, 64, 3, 3] | 36864 |
| encoder.layers.2.block.1.F2.norm.weight | [64] | 64 |
| encoder.layers.2.block.1.F2.norm.bias | [64] | 64 |
| encoder.last.0.F2.conv.weight | [32, 128, 3, 3] | 36864 |
| encoder.last.0.F2.norm.weight | [128] | 128 |
| encoder.last.0.F2.norm.bias | [128] | 128 |
| encoder.last.1.F2.conv.weight | [128, 128, 3, 3] | 147456 |
| encoder.last.1.F2.norm.weight | [128] | 128 |
| encoder.last.1.F2.norm.bias | [128] | 128 |
| decoder.layers.0.up.weight | [4, 32, 4] | 512 |
| decoder.layers.0.block.0.coupling.F.conv.weight | [32, 32, 3, 3] | 9216 |
| decoder.layers.0.block.0.coupling.F.norm.weight | [32] | 32 |
| decoder.layers.0.block.0.coupling.F.norm.bias | [32] | 32 |
| decoder.layers.0.block.1.coupling.F.conv.weight | [32, 32, 3, 3] | 9216 |
| decoder.layers.0.block.1.coupling.F.norm.weight | [32] | 32 |
| decoder.layers.0.block.1.coupling.F.norm.bias | [32] | 32 |
| decoder.layers.1.up.weight | [4, 16, 4] | 256 |
| decoder.layers.1.block.0.coupling.F.conv.weight | [16, 16, 3, 3] | 2304 |
| decoder.layers.1.block.0.coupling.F.norm.weight | [16] | 16 |
| decoder.layers.1.block.0.coupling.F.norm.bias | [16] | 16 |
| decoder.layers.1.block.1.coupling.F.conv.weight | [16, 16, 3, 3] | 2304 |
| decoder.layers.1.block.1.coupling.F.norm.weight | [16] | 16 |
| decoder.layers.1.block.1.coupling.F.norm.bias | [16] | 16 |
| decoder.layers.2.up.weight | [4, 8, 4] | 128 |
| decoder.layers.2.block.0.coupling.F.conv.weight | [8, 8, 3, 3] | 576 |
| decoder.layers.2.block.0.coupling.F.norm.weight | [8] | 8 |
| decoder.layers.2.block.0.coupling.F.norm.bias | [8] | 8 |
| decoder.layers.2.block.1.coupling.F.conv.weight | [8, 8, 3, 3] | 576 |
| decoder.layers.2.block.1.coupling.F.norm.weight | [8] | 8 |
| decoder.layers.2.block.1.coupling.F.norm.bias | [8] | 8 |

**Fig. 7.** Model description of the Chimeric U-Net used in this work.

| Modules | In x Out x H x W | Parameters |
|---|---|---|
| up.weight | [4, 16, 3, 3] | 576 |
| up.bias | [16] | 16 |
| down.weight | [16, 4, 3, 3] | 576 |
| down.bias | [4] | 4 |
| encoder.layers.0.block.0.F2.conv.weight | [16, 16, 3, 3] | 2304 |
| encoder.layers.0.block.0.F2.norm.weight | [16] | 16 |
| encoder.layers.0.block.0.F2.norm.bias | [16] | 16 |
| encoder.layers.0.block.1.F2.conv.weight | [16, 16, 3, 3] | 2304 |
| encoder.layers.0.block.1.F2.norm.weight | [16] | 16 |
| encoder.layers.0.block.1.F2.norm.bias | [16] | 16 |
| encoder.layers.1.block.0.F2.conv.weight | [8, 32, 3, 3] | 2304 |
| encoder.layers.1.block.0.F2.norm.weight | [32] | 32 |
| encoder.layers.1.block.0.F2.norm.bias | [32] | 32 |
| encoder.layers.1.block.1.F2.conv.weight | [32, 32, 3, 3] | 9216 |
| encoder.layers.1.block.1.F2.norm.weight | [32] | 32 |
| encoder.layers.1.block.1.F2.norm.bias | [32] | 32 |
| encoder.layers.2.block.0.F2.conv.weight | [16, 64, 3, 3] | 9216 |
| encoder.layers.2.block.0.F2.norm.weight | [64] | 64 |
| encoder.layers.2.block.0.F2.norm.bias | [64] | 64 |
| encoder.layers.2.block.1.F2.conv.weight | [64, 64, 3, 3] | 36864 |
| encoder.layers.2.block.1.F2.norm.weight | [64] | 64 |
| encoder.layers.2.block.1.F2.norm.bias | [64] | 64 |
| encoder.last.0.F2.conv.weight | [32, 128, 3, 3] | 36864 |
| encoder.last.0.F2.norm.weight | [128] | 128 |
| encoder.last.0.F2.norm.bias | [128] | 128 |
| encoder.last.1.F2.conv.weight | [128, 128, 3, 3] | 147456 |
| encoder.last.1.F2.norm.weight | [128] | 128 |
| encoder.last.1.F2.norm.bias | [128] | 128 |
| decoder.layers.0.up.weight | [32, 128, 2, 2] | 16384 |
| decoder.layers.0.block.0.F2.conv.weight | [64, 64, 3, 3] | 36864 |
| decoder.layers.0.block.0.F2.norm.weight | [64] | 64 |
| decoder.layers.0.block.0.F2.norm.bias | [64] | 64 |
| decoder.layers.0.block.1.F2.conv.weight | [64, 64, 3, 3] | 36864 |
| decoder.layers.0.block.1.F2.norm.weight | [64] | 64 |
| decoder.layers.0.block.1.F2.norm.bias | [64] | 64 |
| decoder.layers.1.up.weight | [16, 64, 2, 2] | 4096 |
| decoder.layers.1.block.0.F2.conv.weight | [32, 32, 3, 3] | 9216 |
| decoder.layers.1.block.0.F2.norm.weight | [32] | 32 |
| decoder.layers.1.block.0.F2.norm.bias | [32] | 32 |
| decoder.layers.1.block.1.F2.conv.weight | [32, 32, 3, 3] | 9216 |
| decoder.layers.1.block.1.F2.norm.weight | [32] | 32 |
| decoder.layers.1.block.1.F2.norm.bias | [32] | 32 |
| decoder.layers.2.up.weight | [8, 32, 2, 2] | 1024 |
| decoder.layers.2.block.0.F2.conv.weight | [16, 16, 3, 3] | 2304 |
| decoder.layers.2.block.0.F2.norm.weight | [16] | 16 |
| decoder.layers.2.block.0.F2.norm.bias | [16] | 16 |
| decoder.layers.2.block.1.F2.conv.weight | [16, 16, 3, 3] | 2304 |
| decoder.layers.2.block.1.F2.norm.weight | [16] | 16 |
| decoder.layers.2.block.1.F2.norm.bias | [16] | 16 |

**Fig. 8.** Model description of the Standard U-Net used in this work.

| Modules | In x Out x H x W | Parameters |
|---|---|---|
| encoder.layers.0.block.0.coupling.F.conv.weight | [8, 8, 3, 3] | 576 |
| encoder.layers.0.block.0.coupling.F.norm.weight | [8] | 8 |
| encoder.layers.0.block.0.coupling.F.norm.bias | [8] | 8 |
| encoder.layers.0.block.1.coupling.F.conv.weight | [8, 8, 3, 3] | 576 |
| encoder.layers.0.block.1.coupling.F.norm.weight | [8] | 8 |
| encoder.layers.0.block.1.coupling.F.norm.bias | [8] | 8 |
| encoder.layers.0.block.2.coupling.F.conv.weight | [8, 8, 3, 3] | 576 |
| encoder.layers.0.block.2.coupling.F.norm.weight | [8] | 8 |
| encoder.layers.0.block.2.coupling.F.norm.bias | [8] | 8 |
| encoder.layers.0.block.3.coupling.F.conv.weight | [8, 8, 3, 3] | 576 |
| encoder.layers.0.block.3.coupling.F.norm.weight | [8] | 8 |
| encoder.layers.0.block.3.coupling.F.norm.bias | [8] | 8 |
| encoder.layers.0.down.weight | [4, 8, 4] | 128 |
| encoder.layers.1.block.0.coupling.F.conv.weight | [16, 16, 3, 3] | 2304 |
| encoder.layers.1.block.0.coupling.F.norm.weight | [16] | 16 |
| encoder.layers.1.block.0.coupling.F.norm.bias | [16] | 16 |
| encoder.layers.1.block.1.coupling.F.conv.weight | [16, 16, 3, 3] | 2304 |
| encoder.layers.1.block.1.coupling.F.norm.weight | [16] | 16 |
| encoder.layers.1.block.1.coupling.F.norm.bias | [16] | 16 |
| encoder.layers.1.block.2.coupling.F.conv.weight | [16, 16, 3, 3] | 2304 |
| encoder.layers.1.block.2.coupling.F.norm.weight | [16] | 16 |
| encoder.layers.1.block.2.coupling.F.norm.bias | [16] | 16 |
| encoder.layers.1.block.3.coupling.F.conv.weight | [16, 16, 3, 3] | 2304 |
| encoder.layers.1.block.3.coupling.F.norm.weight | [16] | 16 |
| encoder.layers.1.block.3.coupling.F.norm.bias | [16] | 16 |
| encoder.layers.1.down.weight | [4, 16, 4] | 256 |
| encoder.layers.2.block.0.coupling.F.conv.weight | [32, 32, 3, 3] | 9216 |
| encoder.layers.2.block.0.coupling.F.norm.weight | [32] | 32 |
| encoder.layers.2.block.0.coupling.F.norm.bias | [32] | 32 |
| encoder.layers.2.block.1.coupling.F.conv.weight | [32, 32, 3, 3] | 9216 |
| encoder.layers.2.block.1.coupling.F.norm.weight | [32] | 32 |
| encoder.layers.2.block.1.coupling.F.norm.bias | [32] | 32 |
| encoder.layers.2.block.2.coupling.F.conv.weight | [32, 32, 3, 3] | 9216 |
| encoder.layers.2.block.2.coupling.F.norm.weight | [32] | 32 |
| encoder.layers.2.block.2.coupling.F.norm.bias | [32] | 32 |
| encoder.layers.2.block.3.coupling.F.conv.weight | [32, 32, 3, 3] | 9216 |
| encoder.layers.2.block.3.coupling.F.norm.weight | [32] | 32 |
| encoder.layers.2.block.3.coupling.F.norm.bias | [32] | 32 |
| encoder.layers.2.down.weight | [4, 32, 4] | 512 |
| encoder.last.0.coupling.F.conv.weight | [64, 64, 3, 3] | 36864 |
| encoder.last.0.coupling.F.norm.weight | [64] | 64 |
| encoder.last.0.coupling.F.norm.bias | [64] | 64 |
| encoder.last.1.coupling.F.conv.weight | [64, 64, 3, 3] | 36864 |
| encoder.last.1.coupling.F.norm.weight | [64] | 64 |
| encoder.last.1.coupling.F.norm.bias | [64] | 64 |
| encoder.last.2.coupling.F.conv.weight | [64, 64, 3, 3] | 36864 |
| encoder.last.2.coupling.F.norm.weight | [64] | 64 |
| encoder.last.2.coupling.F.norm.bias | [64] | 64 |
| encoder.last.3.coupling.F.conv.weight | [64, 64, 3, 3] | 36864 |
| encoder.last.3.coupling.F.norm.weight | [64] | 64 |
| encoder.last.3.coupling.F.norm.bias | [64] | 64 |
| decoder.layers.0.up.weight | [4, 32, 4] | 512 |
| decoder.layers.0.block.0.coupling.F.conv.weight | [32, 32, 3, 3] | 9216 |
| decoder.layers.0.block.0.coupling.F.norm.weight | [32] | 32 |
| decoder.layers.0.block.0.coupling.F.norm.bias | [32] | 32 |
| decoder.layers.0.block.1.coupling.F.conv.weight | [32, 32, 3, 3] | 9216 |
| decoder.layers.0.block.1.coupling.F.norm.weight | [32] | 32 |
| decoder.layers.0.block.1.coupling.F.norm.bias | [32] | 32 |
| decoder.layers.0.block.2.coupling.F.conv.weight | [32, 32, 3, 3] | 9216 |
| decoder.layers.0.block.2.coupling.F.norm.weight | [32] | 32 |
| decoder.layers.0.block.2.coupling.F.norm.bias | [32] | 32 |
| decoder.layers.0.block.3.coupling.F.conv.weight | [32, 32, 3, 3] | 9216 |
| decoder.layers.0.block.3.coupling.F.norm.weight | [32] | 32 |
| decoder.layers.0.block.3.coupling.F.norm.bias | [32] | 32 |
| decoder.layers.1.up.weight | [4, 16, 4] | 256 |
| decoder.layers.1.block.0.coupling.F.conv.weight | [16, 16, 3, 3] | 2304 |
| decoder.layers.1.block.0.coupling.F.norm.weight | [16] | 16 |
| decoder.layers.1.block.0.coupling.F.norm.bias | [16] | 16 |
| decoder.layers.1.block.1.coupling.F.conv.weight | [16, 16, 3, 3] | 2304 |
| decoder.layers.1.block.1.coupling.F.norm.weight | [16] | 16 |
| decoder.layers.1.block.1.coupling.F.norm.bias | [16] | 16 |
| decoder.layers.1.block.2.coupling.F.conv.weight | [16, 16, 3, 3] | 2304 |
| decoder.layers.1.block.2.coupling.F.norm.weight | [16] | 16 |
| decoder.layers.1.block.2.coupling.F.norm.bias | [16] | 16 |
| decoder.layers.1.block.3.coupling.F.conv.weight | [16, 16, 3, 3] | 2304 |
| decoder.layers.1.block.3.coupling.F.norm.weight | [16] | 16 |
| decoder.layers.1.block.3.coupling.F.norm.bias | [16] | 16 |
| decoder.layers.2.up.weight | [4, 8, 4] | 128 |
| decoder.layers.2.block.0.coupling.F.conv.weight | [8, 8, 3, 3] | 576 |
| decoder.layers.2.block.0.coupling.F.norm.weight | [8] | 8 |
| decoder.layers.2.block.0.coupling.F.norm.bias | [8] | 8 |
| decoder.layers.2.block.1.coupling.F.conv.weight | [8, 8, 3, 3] | 576 |
| decoder.layers.2.block.1.coupling.F.norm.weight | [8] | 8 |
| decoder.layers.2.block.1.coupling.F.norm.bias | [8] | 8 |
| decoder.layers.2.block.2.coupling.F.conv.weight | [8, 8, 3, 3] | 576 |
| decoder.layers.2.block.2.coupling.F.norm.weight | [8] | 8 |
| decoder.layers.2.block.2.coupling.F.norm.bias | [8] | 8 |
| decoder.layers.2.block.3.coupling.F.conv.weight | [8, 8, 3, 3] | 576 |
| decoder.layers.2.block.3.coupling.F.norm.weight | [8] | 8 |
| decoder.layers.2.block.3.coupling.F.norm.bias | [8] | 8 |

**Fig. 9.** Model description of the Invertible U-Net used in this work.

### Supplementary Note 8: Bibliography

1. Alexander Buslaev, Vladimir I. Iglovikov, Eugene Khvedchenya, Alex Parinov, Mikhail Druzhinin, and Alexandr A. Kalinin. Albumentations: Fast and flexible image augmentations. *Information*, 11(2), 2020. ISSN 2078-2489. doi: 10.3390/info11020125.
